# Anthracyclins increase free PUFAs and etherPEs with PUFAs as potential hallmarks of lipid peroxidation and ferroptosis

**DOI:** 10.1101/2021.02.16.431377

**Authors:** David Balgoma, Fredrik Kullenberg, Carlemi Calitz, Maria Kopsida, Femke Heindryckx, Hans Lennernäs, Mikael Hedeland

## Abstract

Metabolic and personalized interventions in cancer treatment require a better under-standing of the relationship between the induction of cell death and metabolism. Consequently, we treated three primary liver cancer cell lines with two anthracyclins (doxorubicin and idarubin) and studied the changes of the lipidome. We found that both anthracyclins in the three cell lines increased the levels of polyunsaturated fatty acids (PUFAs) and alkylacylglycerophosphoethano-lamines (etherPEs) with PUFAs. As PUFAs and alkylacylglycerophospholipids with PUFAs are fundamental in lipid peroxidation during ferroptotic cell death, our results suggests supplementa-tion with PUFAs and/or etherPEs with PUFAs as a potential general adjuvant of anthracyclins. In contrast, neither the markers of de novo lipogenesis nor cholesterol lipids presented the same trend in all cell lines and treatments. In agreement with previous research, this suggests that modulation of the metabolism of cholesterol could be considered a specific adjuvant of anthracyclins depend-ing on the type of tumor and the individual. Finally, we discuss the changes in the lipidome in re-lation to the endoplasmic reticulum stress and the sensitivity to anthracyclins of the different cells. In conclusion, our results suggest that the modulation of different lipid metabolic pathways may be considered for generalized and personalized metabochemotherapies.

## 1. Introduction

All cancers are characterized by an inherent metabolic reprograming that promotes tumorigenesis by facilitating and enabling proliferation, metastasis, and resistance to therapies^1,2^. Therefore, metabolomics and lipidomics play key roles in unravelling the metabolic transformation in cancer^3^ and cancer treatment^4^. Hepatocellular carcinoma (HCC) is one of the most common cancers worldwide and it is known to cause profound modifications in lipid metabolism^5^. Among others, the malignant transformation of hepatocytes dysregulates the de novo lipogenesis^5^, which is hallmarked by the up-regulation of triacylglycerides (TGs) with saturated and monounsaturated fatty acids^6^. This altered lipid metabolism is involved in rapid tumor growth and adaptation to the tumor microenvironment^7^.

Clinicians have used anthracyclins, such as doxorubicin (DOX) and idarubicin (IDA), as chemotherapeutic agents against HCC among other solid tumors for more than five decades. Anthracyclins intercalate into the nucleus and mitochondrial DNA and subsequently inhibit the synthesis of proteins and affect the redox state of the cell. Anthracyclins also act on the mitochondrial electron transport and convert oxygen into reactive oxygen species that may cause mitochondrial dysfunction, change the redox state, and induce lipid peroxidation^8^. However, the interplay between anthracyclins and the lipidome is not fully understood. Different anthracyclins establish different physicochemical interaction with lipid membranes depending on the lipid composition^9,10^. For example, IDA is more lipophilic than DOX (logP 1.9 vs logP 1.3) and 7-8 times more potent in vitro^9,11^. Depending on the lipophilic and amphiphilic nature of the anthracyclins, the lipidome of the cell affects drug internalization, and, consequently, its effect^9^. For example, DOX-resistant MCF-7 cells present an enrichment of glycerophospholipids and cholesterol lipids^12^. The changes in the lipidome may affect the further internalization of the drug and the mechanism of action of the drug to induce cell death, in which endoplasmic reticulum (ER) stress plays a key role. This interplay among anthracyclin uptake, the lipidome, and ER stress may partially explain the different sensitivity to anthracyclins in both cancerous and nontransformed cell (e.g. cardiomyocytes). The sensitivity is conditioned by the mechanism of cell death induced by anthracyclins, which depends on the cell type and drug concentration^13^. Consequently, understanding this interplay anthracyclin/lipidome/cell death might open the possibility to: i) improved diagnosis by biomarkers of the tumor resistance to anthracyclins^14,15^, and ii) proposal of novel treatments that may potentiate and/or complement the metabolic transformations caused by anthracyclins to induce cell-death.

Currently there are few studies about the effect of DOX on the lipidome of cancer cells and even less on the effect of other anthracyclins. For example, no lipidomics studies with IDA have been reported (Web of Science, December 2020). In addition, different studies have reported that lipidic modulation affects ER stress, cell death, and cardiotoxicity induced by anthracyclins^2,16,17^. Consequently, more research about the effect of anthracyclins on the cellular tumor lipidome is needed. In this study, we aimed to characterize the lipidic metabolic transformation of cancer cells after exposure to two different anthracyclins. We focused on finding and discussing common and differential changes in the lipidome to different anthracyclins and cell models in order to discuss the mechanism of cell death, the sensitivity to anthracyclins, and potential metabochemotherapies. Consequently, three different primary liver cancer cell lines (HepG2, Huh7, and SNU449) were treated with DOX or the more lipophilic and potent IDA^9,11^. Subsequently, we investigated the regulation of the different lipids by their fold change, and found that both anthracyclins increased the levels of polyunsaturated fatty acids (PUFAs) and alkylacylglycerophosphoethanolamines (etherPEs) with PUFAs in all cell lines (Scheme 1). In the light of previous research, these lipids are involved in programmed cell death by ferroptosis^18^. The changes in cholesterol lipids were cell-type specific, as they increased in Huh7 and SNU449 cells, but did not present a clear trend in HepG2. Similarly, the glycerolipids of saturated and monounsaturated fatty acids did not present a consistent trend for both treatments in both cell lines. These lipids are of interest because they are markers of the liver-X-receptor (LXRs) and sterol regulatory element building proteins (SREBPs) pathways for de novo lipogenesis^6^. Finally, we discussed the relationship between the lipidome, ER stress, and the sensitivity to anthracyclins. From a broad perspective, our results suggest the possibility of using PUFAs and/or etherPEs with PUFAs as general adjuvant of anthracyclins to build new metabochemotherapy treatments.

**Scheme 1.**
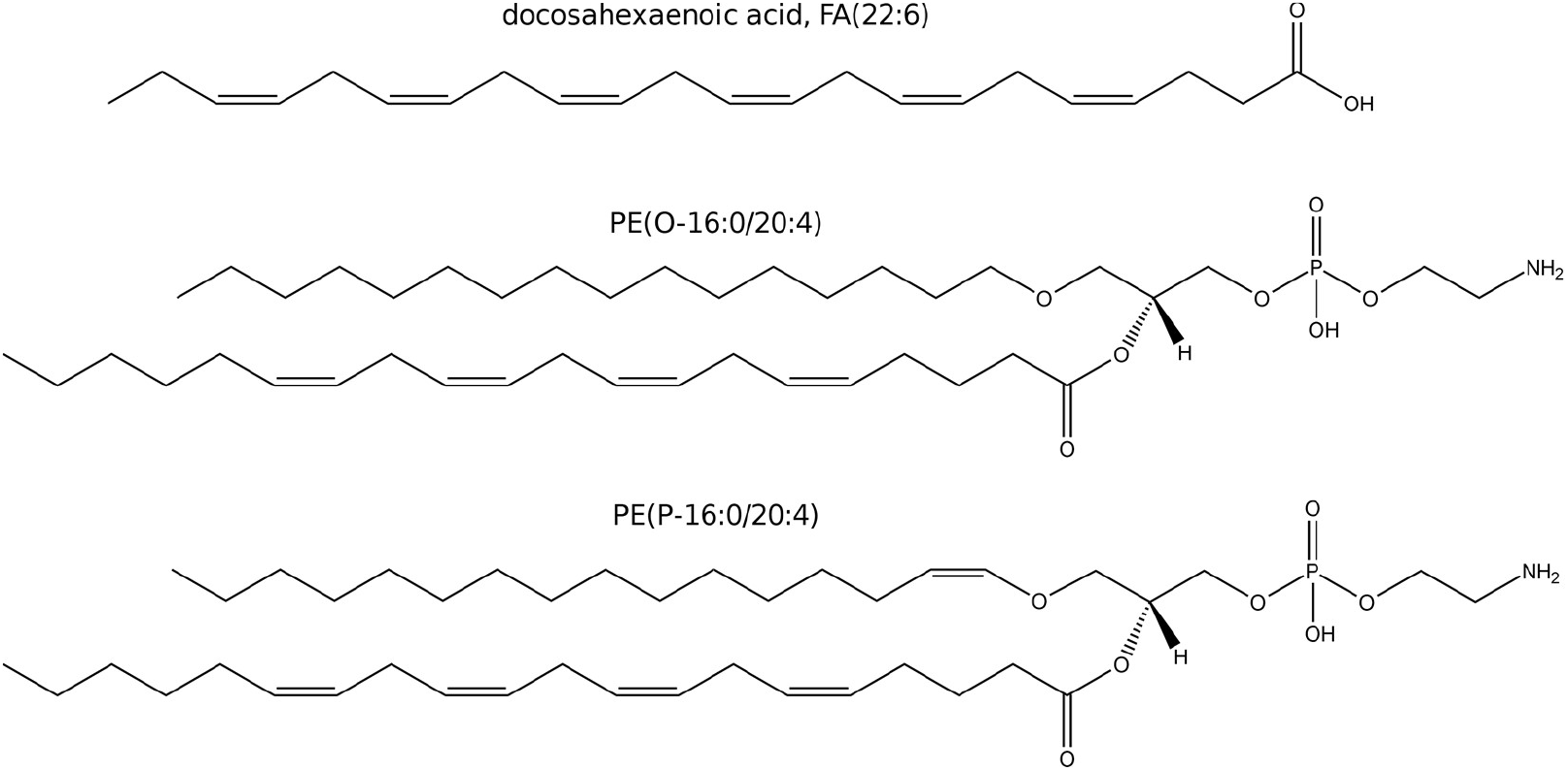
Examples of a PUFA, FA(22:6); and two ether glycerophospholipids: a plasmanyl gly-cerophospholipid of ethanolamine, PE(O-16:0/20:4); a plasmenyl glycerophospholipid of ethanolamine, PE(P-16:0/20:4).

## 2. Materials and Methods

### 2.1. Cell culture and treatment

Three cell lines (HepG2, ATCC^®^ HB-8065™, SNU 449 ATCC^®^ CRL-2234™, and Huh7 from Dilruba Ahmed, Karolinska Institute, Sweden) were cultured at 37°C with 5% CO2 and 95% humidity within a CO2 incubator. HepG2 and Huh7 were cultured in DMEM supplemented with 1% antibiotic antimycotic solution and 10% FBS (cell culture media + FBS: CCMFed). SNU449 was cultured in RPMI medium supplemented with 1% antibiotic antimycotic solution and 10% FBS (cell culture media + FBS: CCMFed). Standard culture medium without supplemented FBS was used during starvation (CCMSM). Misidentification of all cell lines was checked at the Register of Misidentified Cell Lines ^19^. For authentication, extracted DNA from all three cell lines were sent to Eurofins Genomics (Ebersberg, Germany) for cell line authentication using DNA and short tandem repeat-profiles. Mycoplasma contamination was also tested.

Cells were seeded at a density of 4o10^6^ cells per T75 flask (75 cm^2^, 60 ml), and allowed to attach overnight. Prior to treatment, CCMFed was removed and the cells washed with 10 mL of PBS. To allow stabilization of the cell cycle, 15 ml CCMSM was added to each flask containing cells 2 hours prior treatment. Cells were treated for 48 hours with either DOX or IDA in the form of a stock solution in DMSO containing 100 mM anthracycline to achieve the concentrations in Table 1. For the respective controls, the amount of DMSO was matched as for the anthracyclin treatment (vehicle controls). In all cases, the percentage of DMSO was between 0.03% and 0.0005%, which was at least 30 times below the limit at which DMSO presented cytotoxicity in our previous experiments (1% of DMSO, Kullenberg et al., submitted^20^). The exposure concentration of the anthracyclins was chosen to have cell mortality higher than 50% at 48h of treatment. After 48 hours, the cells were washed twice with 5 ml phosphate-buffered saline (PBS). A volume of 5 ml PBS was added to each flask and the cells gently scraped using a cell scraper (TPP). The resulting cell suspension was collected and the cells counted using a TC20™ Automated cell counter and counting slides (Bio Rad). The cell suspension was subsequently centrifuged at 140 g for 5 minutes, the supernatant removed and the pellet was resuspended in 250 μl ice-cold MilliQ water and kept at −80°C until analysis.

**Table 1.** Concentration in cell culture media of DOX and IDA treatments by cell type

| Cell line | DOX concentration (μM) | IDA concentration (μM) |
| --- | --- | --- |
| HepG2 | 0.5 | 0.1 |
| Huh7 | 5 | 0.1 |
| SNU449 | 30 | 1 |

For ER stress imaging and measurement, cells were fixed for 10 minutes in 4% paraformaldehyde and stored at 4°C. Paraformaldehyde fixed cells were washed with tris-buffer saline (TBS) and antigen retrieval was done at 95°C in sodium citrate buffer for 45 minutes. Blocking was followed by an overnight incubation at 4°C with primer antibodies. 40-minute incubation was used for the secondary antibody (Rabbit anti-mouse Alexa Fluor-488 or donkey anti-rabbit Alexa Fluor-633) and cell nuclei were stained with Hoechst for 5 minutes. Images were taken using an inverted confocal microscope (LSM 700, Zeiss) using Plan-Apochromat 20× objectives and the Zen 2009 software (Zeiss). The different channels of immunofluorescent images were merged using ImageJ software. Quantifications were done blindly with ImageJ software by conversion to binary images for each channel and automated detection of staining on thresholded images.

### 2.2. Lipidomics analysis and data pre-treatment

After randomization of the order of sample preparation, 100 μL of acetonitrile/isopropanol 50:50 were added to 10 μL of sample. Proteins were precipitated overnight at −20°C and the supernatant was recovered after centrifugation (18000 RCF, 10 min). A quality control sample was built by pooling 10 μL from every extract. The samples were analyzed as in Balgoma et al.^21^. The samples were injected in a randomized way on an Acquity UPLC hyphenated to a Synapt G2 Q-ToF (Waters) with electrospray ionization. The samples were analyzed in both positive and negative modes. The quality control pool was injected every five injections of samples. Lipids were identified as in Balgoma et al.^21^ by the m/z of their adducts and their patterns of fragmentation. Due to the number of lipids, only the key species are represented in the main text. We detected 451 species of lipids peaks corresponding to free fatty acids (FA), lysophosphatidylcholine (LPC), phosphatidylcholine (PC), alkylacylglycerophospholipids of choline (etherPC), lysophosphatidylethanolamine (LPE), phosphatidylethanolamine (PE), etherPE, phosphatidylglycerol (PG), lysophosphatidylinositol (LPI), phosphatidylinositol (PI), lysophosphatidylserine (LPS), phosphatidylserine (PS), diacylglycerols (DG), TG, free sterols, cholesteryl esters (CE), ceramides (Cer) and sphingomyelin (SM).

### 2.3. Data analysis

We addressed the concerns about using p-values and “statistical significant” discoveries exposed by researchers, statisticians, and the American Statistical Association^22–24^. Consequently, we analyzed the changes in the lipidome by using fold changes (with confidence intervals) between the treatments and the control^25^.

The cell size was quantified by the weighted mean of diameter of cells. The variability was calculated by the standard error of the mean of the three replicates.

Regarding lipidomics data, the areas of the peaks were normalized by the number of cells extracted. The changes in the lipidome were studied by the relative changes (increase/decrease) between the cells treated with anthracyclins and the cells treated with their respective controls (vehicle). These relative changes were quantified by the log(fold change), i.e. the natural logarithm of the ratio of the average of the normalized signal of the treated group divided by the average of the normalized signal of the control group (vehicle). The confidence interval of the log(fold change) was determined by the 10,000 bootstrap resampling simulations. The limits of the intervals of confidence were selected by the percentiles 2.5% and 97.5% of the simulations; the central measurement was characterized by the mean of the simulations. The values under the limit of detection (left-censored) were imputed by fitting the peak areas of a lipid to a normal distribution and random sampling of this distribution below the minimum detected signal. Fold changes with more than one value under the limit of detection in any group were discarded.

### 2.4. Software and graphical material

Waters’ .Raw data files were transformed into .CDF format by Databride (Masslynx 4.1). Mass spectrometric data were pretreated with packages mzR 2.22.0 and XCMS 3.10.2 in R 4.0.3 “Bunny-Wunnies Freak Out”. Graphical material was generated with R (packages forestplot 1.10 and ggplot2 3.3.2) and processed with Inskape 1.0.1 and Gimp 2.10.22. Microscopy images were obtained in Zen 2009 software (Zeiss) and further quantified and exported using ImageJ software.

## 3. Results

### 3.1. Number and size of the recovered cells

As expected, DOX and IDA induced a strong reduction in cell number when compared with the vehicle control. The respective vehicle controls did not induce cell death nor proliferation (Figure 1). In all cases, anthracyclins induced a reduction in cell number higher than 50%, when compared with the vehicle.

**Figure 1.**
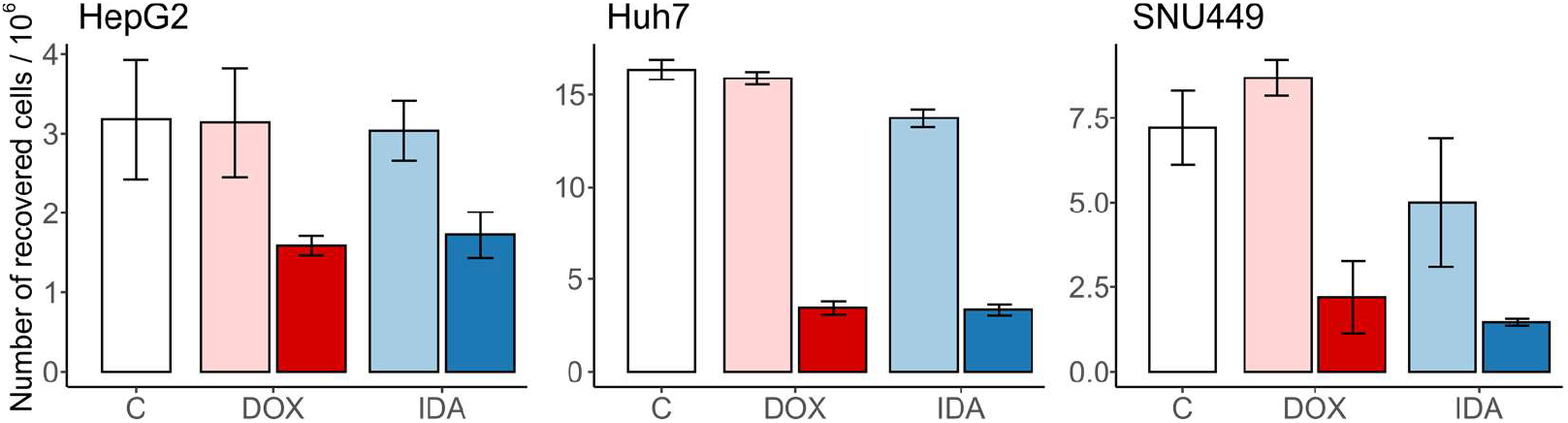
Number of recovered cells after treatment with anthracyclins. The untreated control (C) is in white, the vehicles of DOX and IDA are in light red and blue, respectively. The treatments of DOX and IDA are in dark red and blue, respectively. Whiskers represent the interval for the standard error of the mean.

Regarding the cell size, the vehicle controls did not change the diameter of the cells when compared with the untreated control (Figure 2). The treatment with anthracyclins did not change the cell volume for HepG2 nor Huh7. In contrast, we observed an increased cell diameter in DOX and IDA treated SNU449-cells (Figure 2).

**Figure 2.**
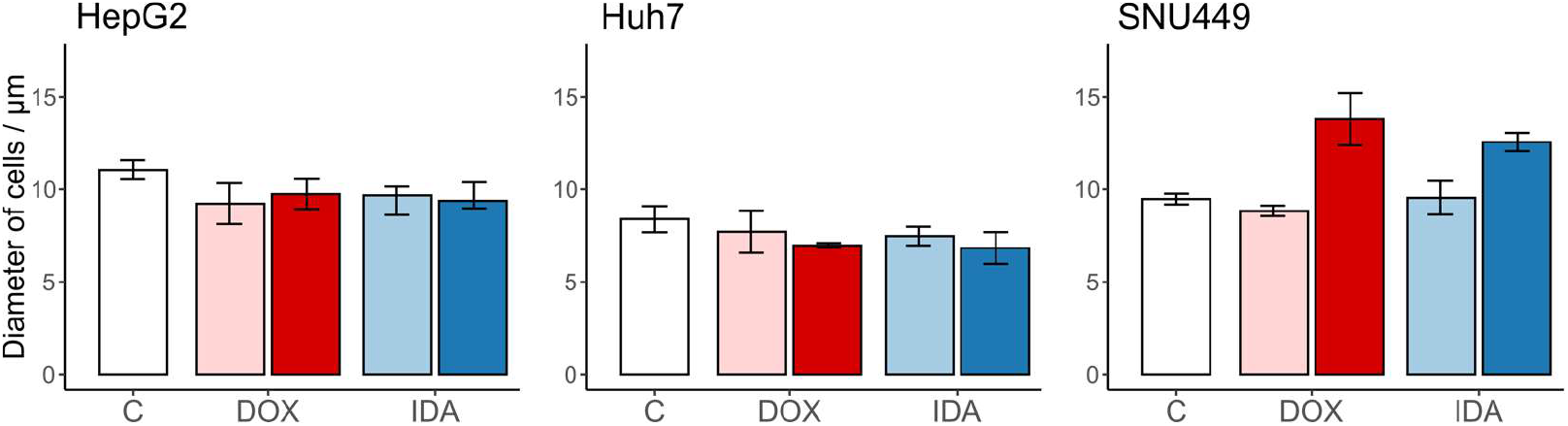
Size of the recovered cells (diameter, μm) after treatment with anthracyclins. The untreated control (C) is in white, the vehicles of DOX and IDA are in light red and blue, respectively. The treatments of DOX and IDA are in dark red and blue, respectively. Whiskers represent the interval for the standard error of the mean.

### 3.2. sAnthracyclins increase etherPEs in all cell types

To estimate the regulation of the total amount of lipids per family, we summed the normalized signal of every lipid family and calculated the logarithm of the fold change with respect to the vehicle-treated cells (Figure 3).

**Figure 3.**
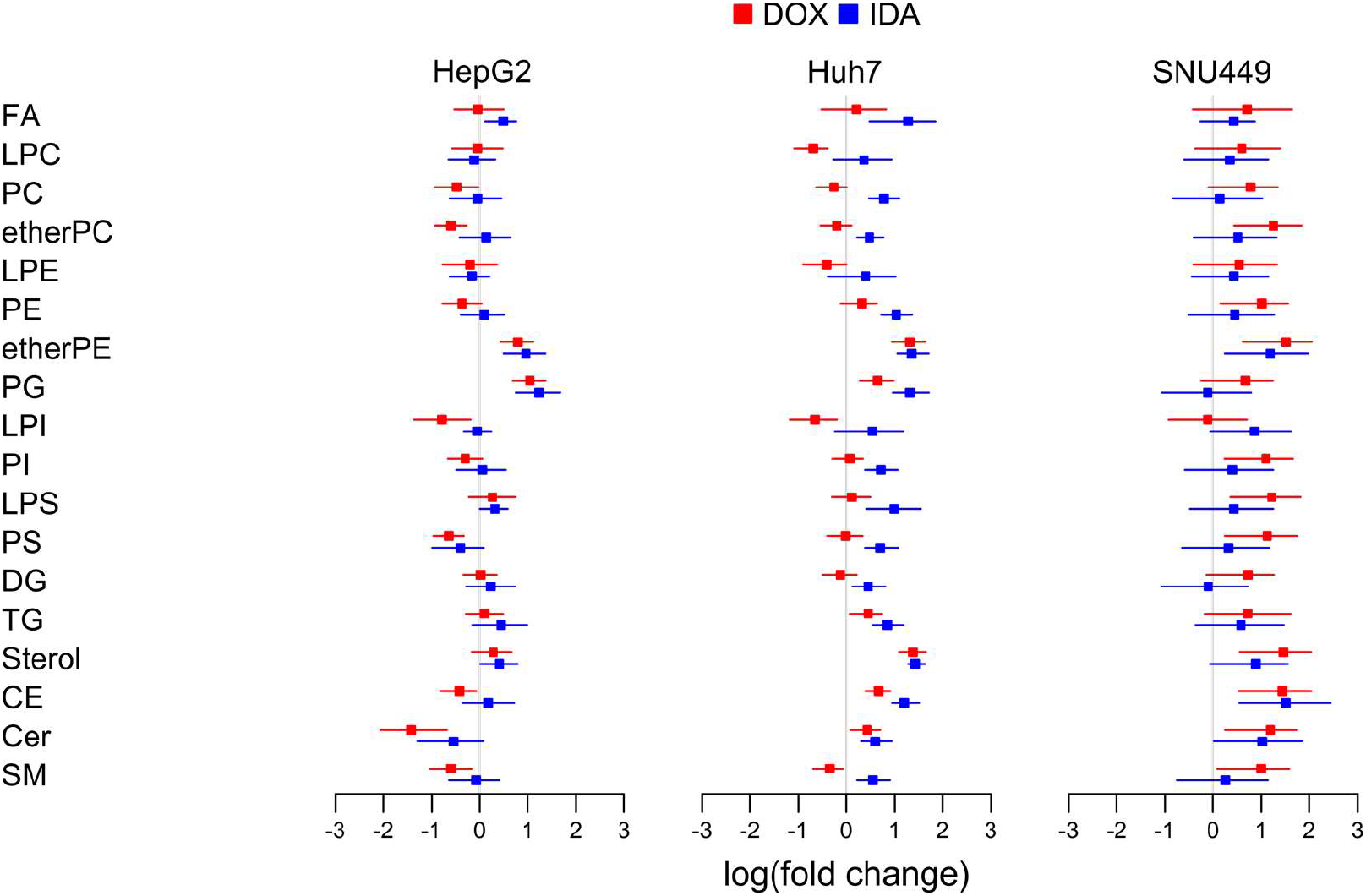
Logarithm of the fold change of the eighteen families of lipids in each cell line. We summed the signal of the lipids per type of cell, treatment, and replicate

In general, the different families of lipids presented no clear trend for all lipids for the three cell types. Regarding the ensemble of the lipid families for every cell type, the lipidome showed a trend to increase in SNU449, especially with DOX (Figure 3).

Regarding common trends for both treatments and the three cell lines, we observed an increase of etherPEs. While most families of lipids increased in SNU449, the fold change of etherPEs was the highest, together with cholesterol lipids. This behavior points to a common and general metabolic effect of anthracyclins in etherPEs beyond the specificities of the metabolism of every cell line.

Regarding disparate trends for the different cell lines, cholesterol lipids (sterols and CEs) presented a trend to increase in Huh7 and SNU449, but not in HepG2. PGs presented a trend to increase in HepG2 and Huh7, but not a clear trend in SNU449. PSs and the sphingolipids (Cer and SM) presented a trend to decrease in HepG2, no clear trend in Huh7, and a trend to increase in SNU449. As highlighted before, the increase in these lipids in SNU449 was in a context of general increase in lipids in this cell line. Furthermore, Huh7 treated with IDA presented a general trend to increase.

### 3.3. Anthracyclins increase all etherPEs with PUFAs, but not etherPC

To study which lipid pathways were responsible for the upregulation of etherPEs in Figure 3, we represented the fold change of the individual species of alkylacylglycerophospholipids (etherGLs) and their fatty acid composition (Figure 4).

We only detected one species of etherPC, PC(O-16:0/18:1). This species did not present a clear trend for all cells and treatments (Figure 4). Regarding etherPEs, we detected six different species and all contained PUFAs. They corresponded to plasmanyl (ether bond) or plasmenyl (vinyl ether bond, Scheme 1), but their mass spectrometric characterization did not allow to distinguish these possibilities from each other. The fold change in etherPEs presented a common trend to increase in the three cell lines with both anthracyclins. We conclude that the upregulation of etherPEs with PUFAs (but not all etherGLs) was a general metabolic trait of anthracyclins.

**Figure 4.**
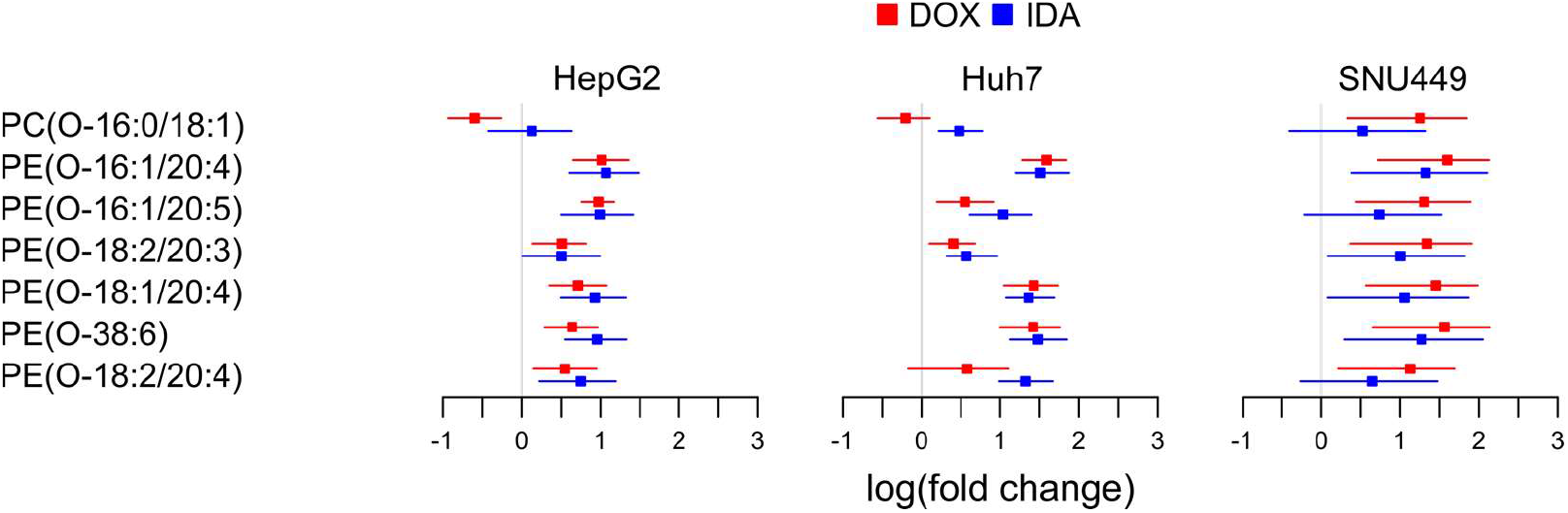
Logarithm of the fold change of the etherGLs. For etherPEs, the structure could also correspond to plasmenyls (e.g. PE(P-16:0/20:4) for PE(O-16:1/20:4))

### 3.4. Anthracyclins increase free PUFAs

To study if the availability of free PUFAs may partially control the increase of etherPEs with PUFAs, we investigated the fold change of FAs.

Despite the different regulation of different fatty acids, FA(20:4) and FA(20:5) showed a clear trend to increase in all cell lines treated with either DOX or IDA (Figure 5). Other PUFAs, such as FA(20:3) and FA(22:6) also presented a trend to increase, but to a lower degree, especially in SNU449. This behavior suggests that free PUFAs are more available in all anthracyclin-treated cells.

**Figure 5.**
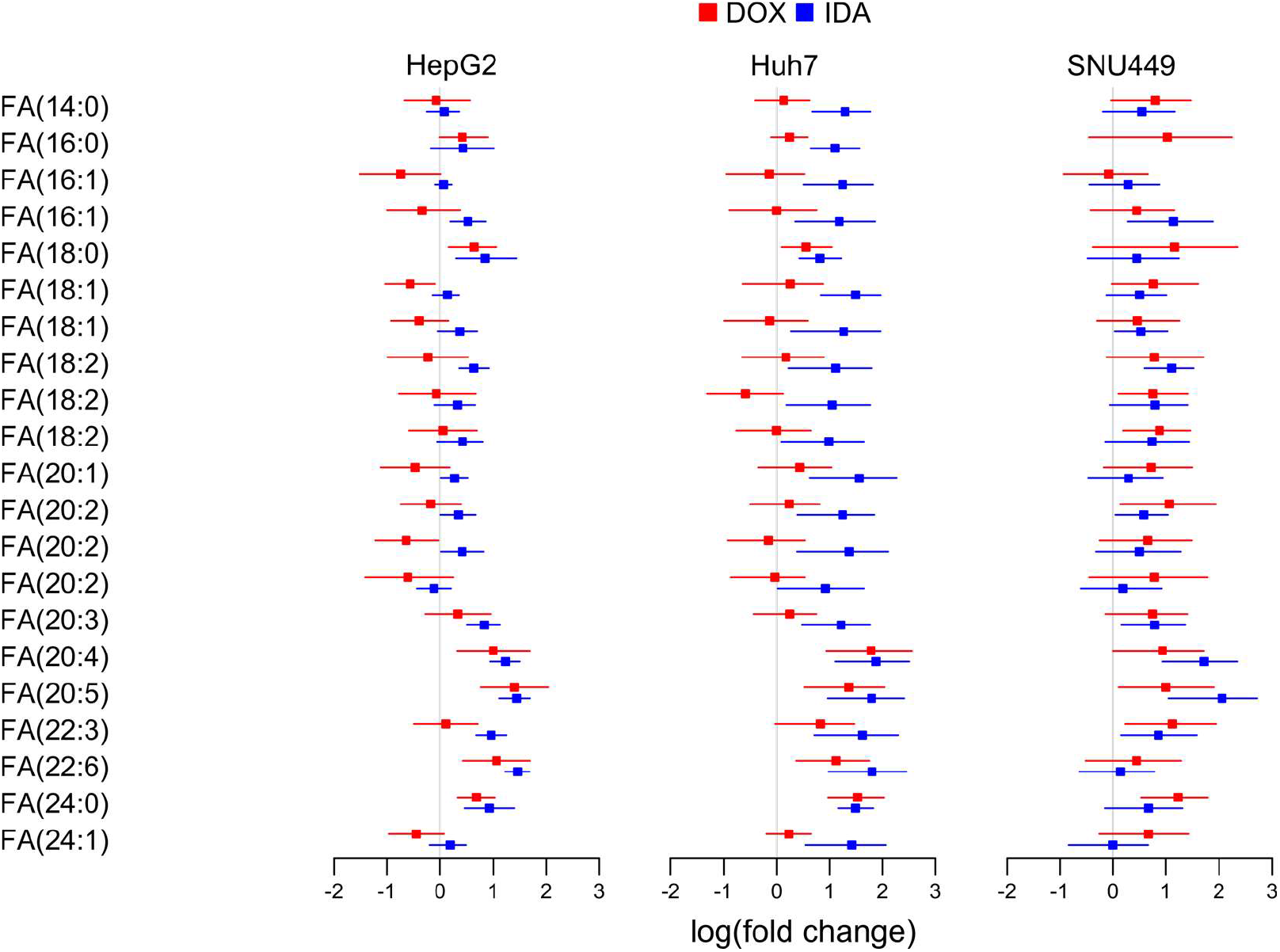
Logarithm of the fold change of free fatty acids (FAs) in each cell line.

### 3.5. Anthracyclins increase other glycerolipids with PUFAs

Radiolabeling experiments have shown that the glycerophospholipids of choline and ethanolamine contain most of PUFAs^26^. Consequently, we studied the levels of glycerolipids with PUFAs to investigate if a net release of PUFAs would explain the increase of free PUFAs. We observed that different molecular species presented different regulation (Figure 6), which might be explained by the different substrate selectivity in the enzymes involved in the liberation and incorporation of PUFAs into glycerolipids ^27^. However, the major species tended to increase in all cells with all treatments. This was the case of PC(16:0/20:4), PC(18:1/20:4), PE(18:0/20:4), PI(18:0/20:4), PC(16:0/20:5), PC(18:0/20:5), and PC(18:1/20:5) (Figure 6). Taken together, these upregulations suggest that the increase of free PUFAs (Figure 5) was not due to a net-release of PUFAs from glycerolipids.

**Figure 6.**
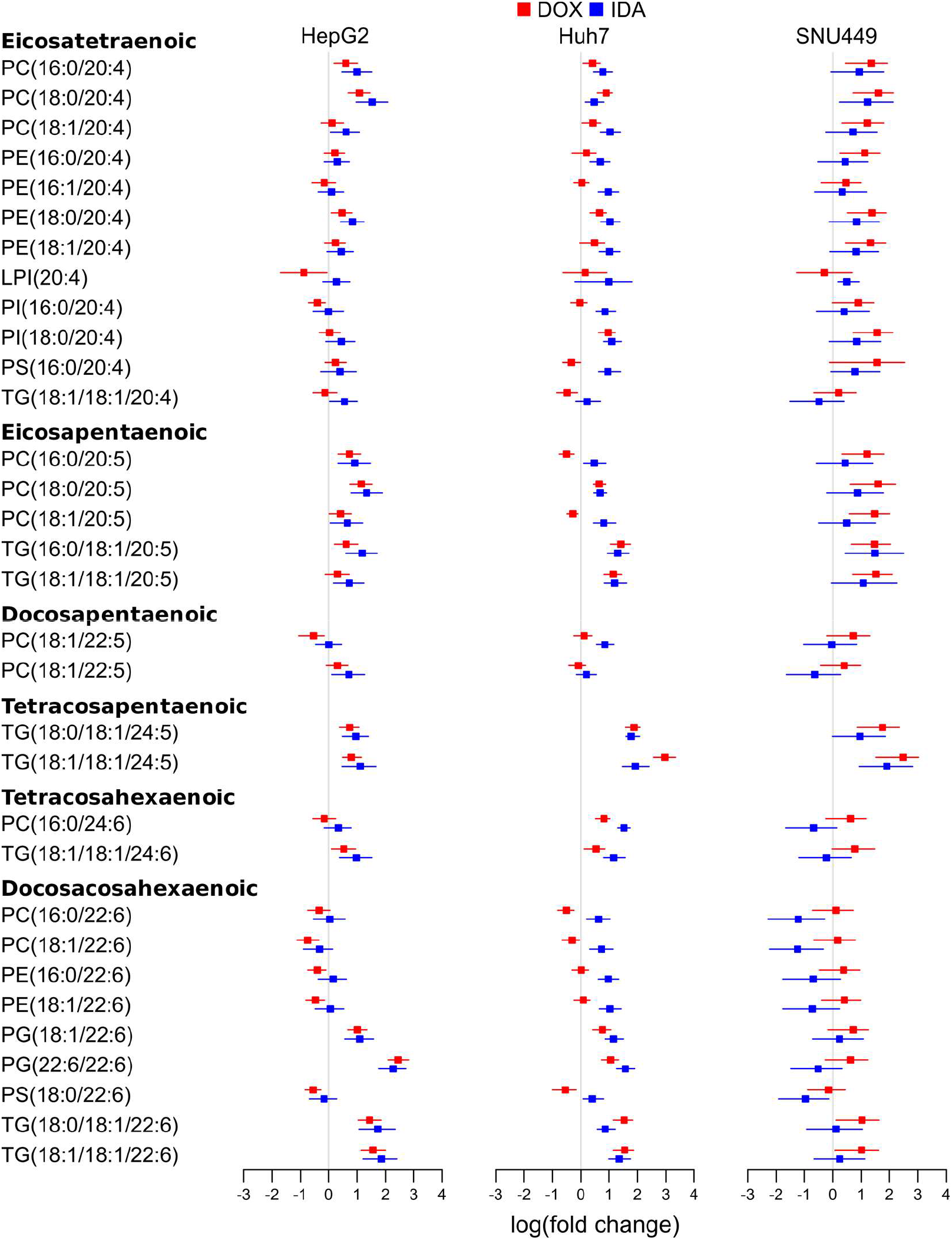
Logarithm of the fold change of selected glycerolipids with PUFAs detected in each cell line.

Interestingly, FA(22:6) (Scheme 1) might present different regulation than other PUFAs, as it is produced from FA(24:6) by peroxisomal β-oxidation^28^. The major species of glycerolipids with FA(22:6) in PC and PE did not present a clear trend to decrease following anthracyclin treatment in any of the cell lines (Figure 6). Some minor species, such as PG(22:6/22:6) and TG(18:0/18:1/22:6) presented a trend to increase in treated HepG2 and Huh7. Nevertheless, these species did not present a clear increase in SNU449. In summary, the data do not suggest a general decrease for all cell lines of neither free nor esterified FA(22:6) (Figures 5 and 6).

### 3.6. Regulation of glycerolipids with saturated and monounsaturated fatty acids

Glycerolipids with FA(16:0), FA(16:1), FA(18:0) and FA(18:1) are markers of LXR and SREBP pathways for de novo lipogenesis^6^. To evaluate if anthracyclins affect the de novo lipogenesis in the three cell lines, we studied the changes of these lipid species (Figure 7).

**Figure 7.**
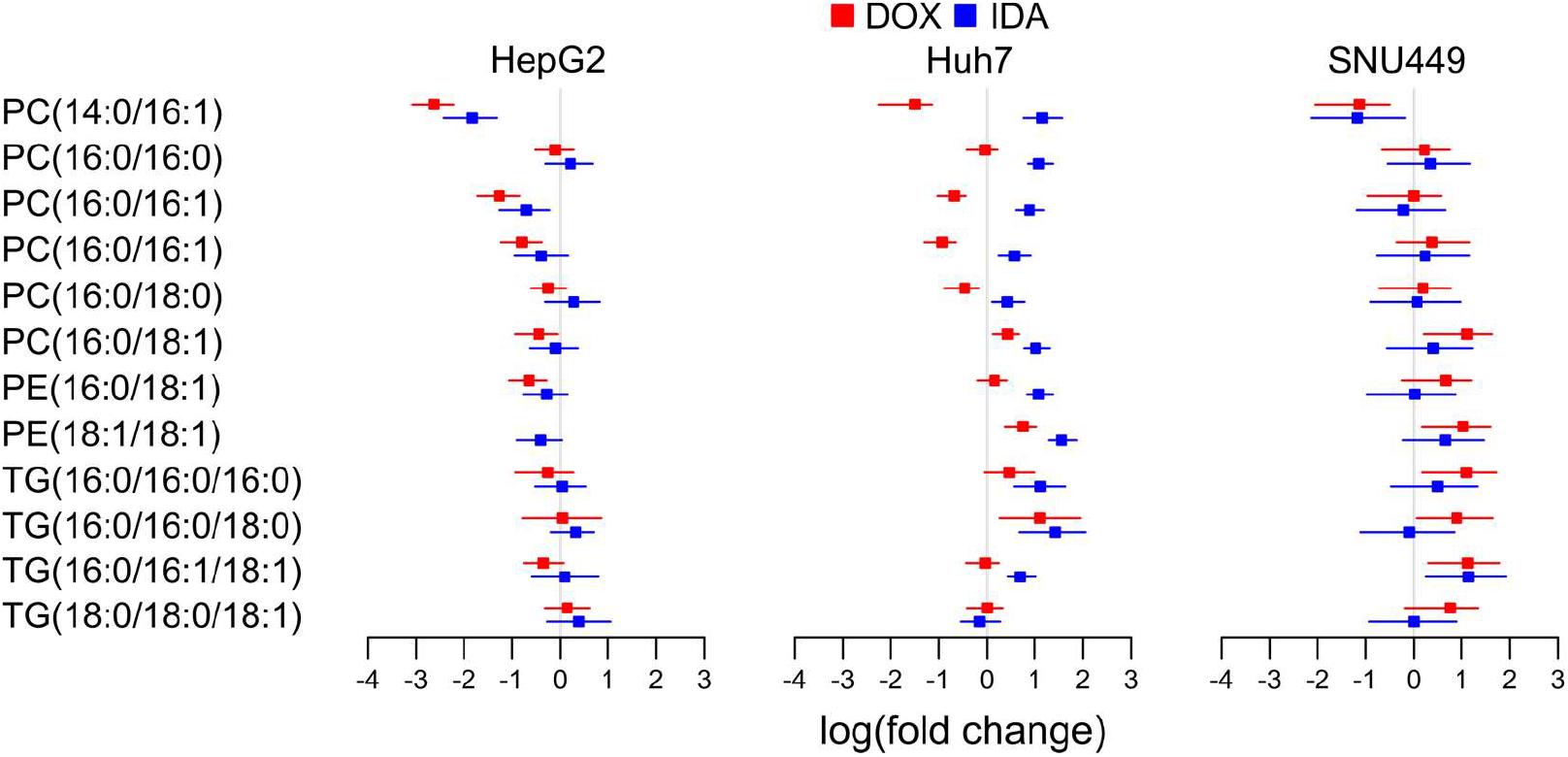
Logarithm of the fold change of selected glycerolipids with saturated and monounsaturated fatty acids in each cell line.

Despite the specific decrease in species like PC(14:0/16:1) in HepG2, the general overview of the main glycerolipid species with saturated and monounsaturated fatty acids did not show any clear trend. Only in Huh7 treated with IDA there was an increase in these species, except for TG(18:0/18:0/18:1). This suggests that anthracyclins do not affect LXRs nor SREBPs de novo lipogenesis in a general way, but it could depend on the cell type-treatment combination.

### 3.7. DOX increases markers of ER stress

To study ER stress, we determined the levels of DNA damage-inducible transcript 3 (a.k.a. C/EBP homologous protein, CHOP) and binding immunoglobulin protein (BIP) through immunocytochemistry (Figure 8). These two markers increased after DOX treatment, which confirms that anthracyclins induce ER stress. The increase was stronger for HepG2 and Huh7, than for SNU449.

**Figure 8.**
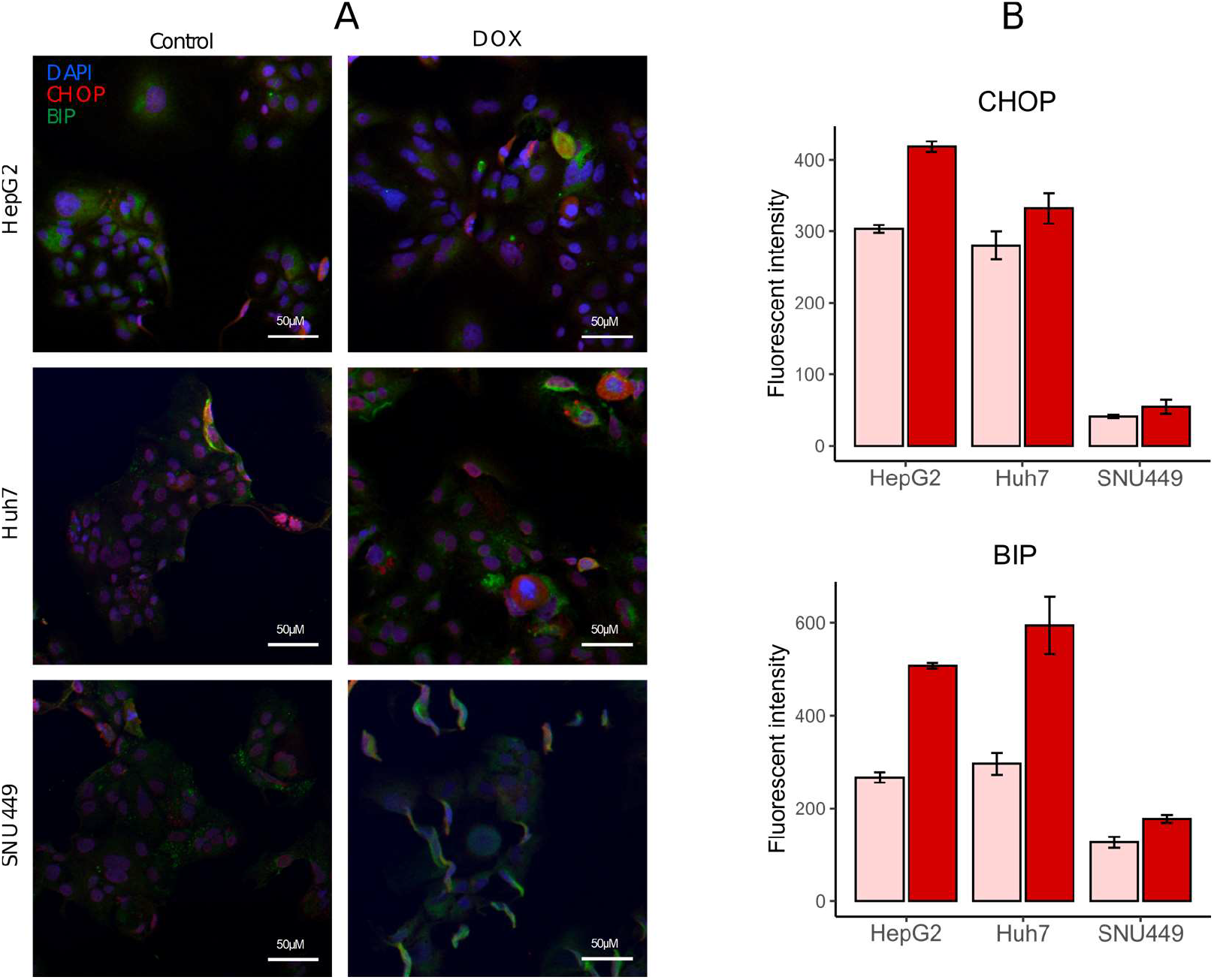
DOX treatment increase markers of ER stress. A) Representative images of HepG2, Huh7, and SNU449 tumor cells stained with antibodies against DAPI (blue), CHOP (red), and BIP (green). B) Quantification of the levels of CHOP and BIP in the cells with vehicle controls (light red) and treatments with DOX (dark red).

## 4. Discussion

Here we present that anthracyclins elicit the upregulation of PUFAs and etherPEs with PUFAs in different primary liver cancer cells. By using DOX and IDA in three different cell lines, we present new evidence that this is a general effect of anthracyclins on the metabolism of cancer cells. We also present evidence that the regulation of cholesterol lipids and de novo lipogenesis by anthracyclins is cell-type dependent for three different cell lines of primary liver cancer. As etherPEs with PUFAs are involved in programmed cell death by ferroptosis^18^ and statins have been suggested as anthracyclin adjuvants^29^, this metabolic characterization may help to design general and personalized metabochemotherapies for improved cancer treatments.

### 4.1. Comparison with other lipidomics studies

We did not observe a general change of lipids in HepG2 and Huh7, except for the increase of PUFAs, etherPEs with PUFAs, and PGs (Figure 3). This observation agrees with the recent study by Newell et al. in 2020, which does not report strong changes in the lipid families in breast cancer cells (MDA-MB-231 and MCF-7) treated with DOX in combination with fatty acids^30^. They did not include etherPEs nor PGs in their study, so, as new result, we observed that both anthracyclins upregulated these families of glycerophospholipids in HepG2 and Huh7 (PUFAs and etherPEs with PUFAs increased in SNU449 as well, Figure 3).

In contrast, Achkar et al. in 2020 found a general downregulation of most metabolites and lipids (including etherPEs with PUFAs) in egg-developed tumors (MDA-MB-231) treated with DOX^31^. This finding is in apparent contradiction to our results and those reported by Newel et al. in 2020^30^. Nevertheless, the study of Achkar et al. in 2020 about DOX-treated solid tumors contained alive, damaged, and dead cell debris^31^. We investigated the change in the lipidome of viable cells and normalized the amount of lipids to the number of cells. The differences in the cell subpopulation and the normalization may explain this observed discrepancy.

In conclusion, our study complements previous studies as we observed that PUFAs and etherPEs with PUFAs were increased in general after the treatment with anthracyclins in three different cell lines. This observation suggests that the increase of these lipids is a general metabolic trait of anthracyclins in liver cancer cells.

### 4.2. Regulation of the de novo lipogenesis of fatty acids, glycerolipids, and cholesterol

LXRs and SREBPs control the expression of the enzymes responsible for the de novo lipogenesis of fatty acids, glycerolipids, and cholesterol^32^. The increase in de novo lipogenesis of fatty acids and glycerolipids is characterized by the upregulation of glycerolipids with saturated and monounsaturated fatty acids^6^. In this context, we did not observe a general up or downregulation of these glycerolipids, which suggests that anthracyclins do not modify the de novo synthesis of fatty acids in general (Figure 7). Regarding the regulation of cholesterol, we observed a trend to upregulation of cholesterol lipids in Huh7 and SNU449 (Figure 3). However, this trend was not clear for HepG2. Consequently, our data do not suggest a general trend for the de novo lipogenesis of neither fatty acids nor cholesterol.

Previous studies also report disparate results regarding the regulation of LXR/SREPB pathways by anthracyclins. On the one hand, it has been reported that DOX slightly upregulates SREBPs in the kidney with disparate behavior of the enzymes responsible for the de novo synthesis of fatty acids and cholesterol^33^. In cardiomyocytes, DOX also upregulates oxysterols and hence elicits LXR activation^34^. On the other hand, other studies report downregulation of 3-hydroxy-3-methyl-glutaryl-coenzyme A reductase (HMGCR) in epidermal and prostate cancer cell lines^35^. Furthermore, chemotherapy may modify the cholesterol efflux from the environment of the cell, which could affect the cholesterol content in many cellular compartments^36^.

Taken our results and previous research together, we conclude that the up or downregulation of the LXR/SREBP-mediated de novo lipogenesis of fatty acids, glycerolipids, and cholesterol was not a general effect of anthracyclins. Nevertheless, as we saw different regulation for three different lines of the same type of cancer, it could depend on the cell type, tumor, or even individual genotype and phenotype.

### 4.3. Regulation of the increase of etherPEs with PUFAs and PUFAs by anthracyclins

The synthesis of etherGLs is initiated in the peroxisome and completed in the ER, where the activity of alcohol reductase FAR1 is the limiting step (Scheme 2)^37–39^. The amount of etherGLs is regulated by: i) the downregulation of FAR1 by the increase of etherGLs in the inner leaflet of the plasma membrane, and ii) the lysoplasmalogenase activity^40^. While we observed an upregulation of etherPEs with PUFAs in all cells and treatments (Figure 4), we detected an etherPC, PC(O-16:0/18:1), that did not present a consistent increase in all treatments and cells. This differential behavior suggests that the upregulation of etherPEs with PUFAs was independent of: i) a putative increase of the synthesis of 1-O-alkyldihydroxyacetone phosphate in the peroxisome (Scheme 2), or ii) a putative decrease in the lysoplasmalogenase activity. Free PUFAs increased in a general way. Consequently, it seems plausible that the availability of PUFAs was the metabolic regulation responsible for the increase of etherPEs with PUFAs.

**Scheme 2.**
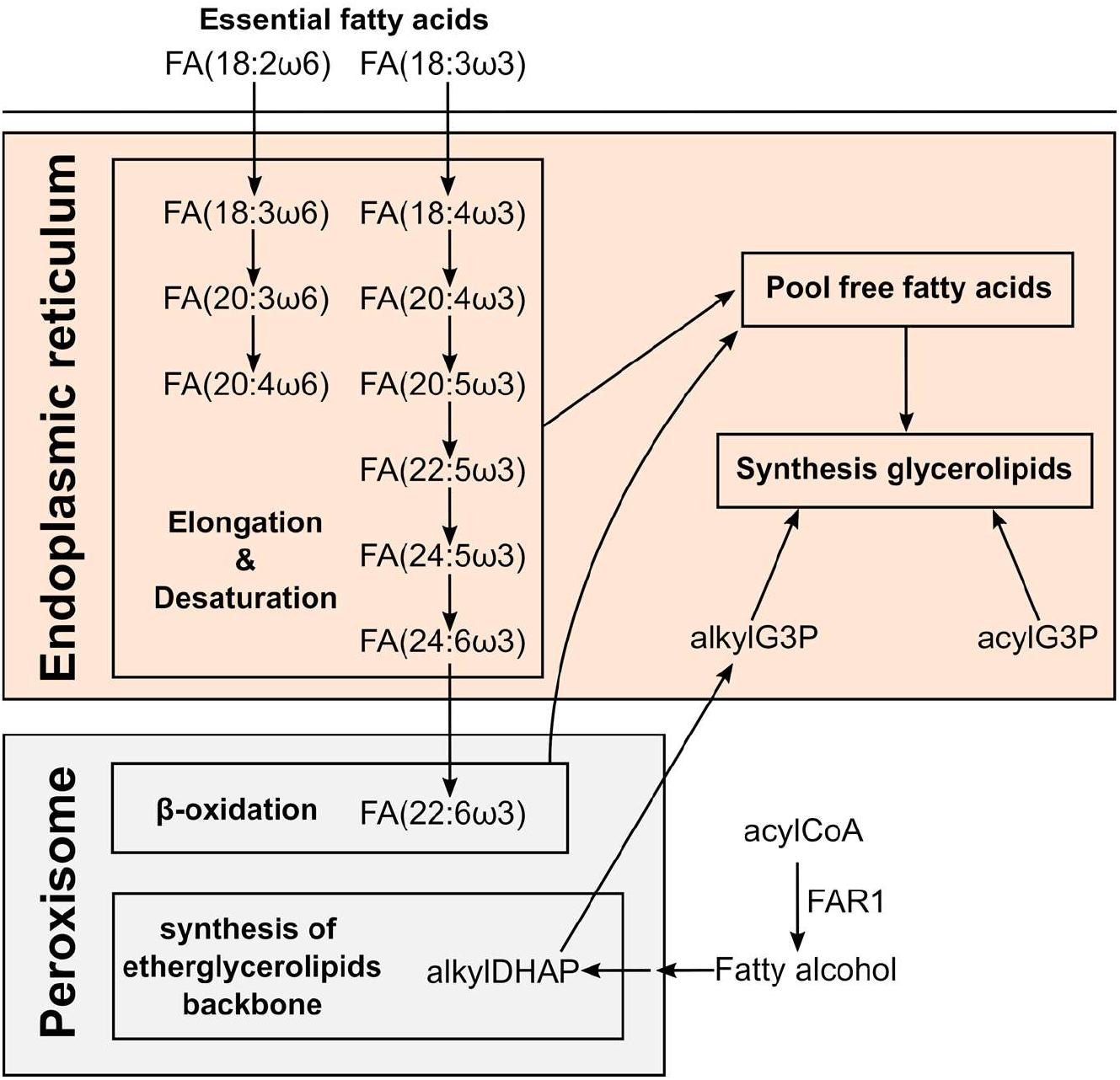
Key molecules in the biosynthetic route of PUFAs and ether glycerolipids in different human subcellular compartments. Adapted from ^41–43^.

The levels of free PUFAs could be increased by three different factors: i) the increase of their synthesis by elongation and desaturation of essential linoleic and linolenic acids^42^; ii) the balance between their liberation from and incorporation into glycerolipids^26,27,44,45^; and/or iii) the downregulation of their degradation by peroxisomal β-oxidation^37^. Consequently, we analyzed the levels of PUFAs in different glycerolipids to discuss the potential involvement of these three processes. On the one hand, the general trend to upregulation of FA(20:4) and FA(20:5) in glycerolipids (Figure 6) suggests that their net-release is not responsible for the increase of their free form. On the other hand, their degradation is initiated in the peroxisome by β-oxidation. Interestingly, docosahexaenoic acid, FA(22:6), is not synthesized by elongation and desaturation in the ER, but by β-oxidation of FA(24:6) in the peroxisome (Scheme 2)^46^. Consequently, the decrease of free and esterified FA(22:6) would be indicative of the downregulation of peroxisomal β-oxidation and, consequently, of the degradation of PUFAs. However, we did not observe a general downregulation of FA(22:6) neither free nor esterified. In fact, we rather observed a trend to increase in specific lipid species in HepG2 and Huh7 (Figure 6). The behavior of FA(22:6) suggests that neither the activity of peroxisomal β-oxidation nor the degradation of PUFAs were downregulated as a general characteristic of anthracyclins.

We conclude that the observation of the ensemble of free PUFAs and glycerolipids with PUFAs suggests the upregulation of the elongation and desaturation of linoleic and linolenic acids as responsible for the increase of free PUFAs and etherPEs with PUFAs (Scheme 2).

### 4.4. Role of PUFAS and etherPEs with PUFAs: potential hallmark of programmed cell death by ferroptosis

Two main lines of research have related PUFAs and cell-death induced by anthracyclins: i) PUFAs enhance the effect of DOX on cell death, and ii) DOX induces ferroptosis, in which PUFAs, etherPEs with PUFAs, and ER stress are involved.

For more than 25 years, it has been known that extracellularly added PUFAs enhance the sensitivity of tumor cell lines to DOX^47,48^. Among different PUFAs, FA(22:6) presents the highest enhancement of cytotoxicity^48^. This effect has also been demonstrated in animal models^30^. However, the mechanism(s) of how PUFAs enhance anthracyclin-induced cell death is(are) not fully understood. Pioneering studies found that PUFA peroxidation might play a key role in boosting the oxidative stress in the tumor cell^49^. In this context, we observed that anthracyclins alone elicited the upregulation of PUFAs. Interestingly, supplementation of cells with PUFAs also enhances cell death by ferroptosis, the other line of research^50^. Considering that we observed a general increase of PUFAs and that DOX-induced cell death has been associated with ferroptosis^51^, it seems plausible that this line of research converges with ferroptosis research. Consequently, the enhancement of cell death induced by PUFAs might be explained by boosting lipid peroxidation in anthracyclin-induced ferroptosis^52^.

Ferroptosis, a term for a non-apoptotic cell death introduced in 2012, is characterized by the intracellular accumulation of lipid hydroperoxides^50,53^. Lipid peroxides yield aldehydes that react with intracellular and membrane proteins, compromising their function. To achieve lipid peroxidation, ferroptosis requires changes in a plethora of pathways, such as iron, glutaminolysis, and cholesterol and mevalonate pathways^53^. Furthermore, ferroptosis does not imply modifications in the morphology of the cell, except for the mitochondria^54^.

In the context of lipid peroxidation and ferroptosis, first, our analysis about the regulation of PUFAs indicates that their increase was due to the upregulation of the elongation and desaturation (Section 4.3, Scheme 2). In this process, Δ6-fatty acid desaturase 2 (FADS2) is necessary for the synthesis of PUFAs^55^. Interestingly, the knockdown of FADS2 presents attenuated lipid peroxidation in Huh7 cells^56^. Consequently, the upregulated elongation and desaturation to yield PUFAs (Section 4.3) is consistent with the increase in lipid peroxidation and ferroptosis.

Second, we found a general increase of etherPEs with PUFAs. It is known that: i) glycerophospholipids of ethanolamine with PUFAs are necessary for ferroptosis^57^; and ii) etherGLs with PUFAs present a critical contribution to ferroptosis^18^. It is plausible that the increase of etherPEs with PUFAs can be considered a hallmark of ferroptosis in primary liver cancer treated with anthracyclins. The reason why etherGLs with PUFAs are important in ferroptosis is not clear. Interestingly, they localize in the internal leaflet of the plasma membrane^40^, which suggests that lipid peroxidation in this region of the cell plays a key role in the cell death by ferroptosis.

Finally, regarding the morphology, we found that HepG2 and Huh7 did not change their size upon anthracyclin treatment. However, SNU449 increased in size with both DOX and IDA. Furthermore, we observed an increase in PUFAs and etherPEs with PUFAs in SNU449, but in the context of a general increase of lipids (Figure 3). EtherPEs with PUFAs presented a slightly higher fold change than other lipids in SNU449 (Figure 3), but the overall change of the lipidome and the increase in size suggest that the enrichment of lipids with PUFAs was diluted in the membranes. Consequently, it seems plausible that lipid peroxidation in SNU449 was quenched.

Considering the discussion in the three previous paragraphs, we conclude that the changes in the lipidome and the cell size suggest that ferroptosis may drive the cell death for HepG2 and Huh7. However, other mechanisms apart from lipid peroxidation and ferroptosis may drive the cell death for SNU449. This suggestion is consistent with previous studies, as the mechanism of cell death depends on the concentration of anthracyclins and the cell type^13^.

### 4.5. Interplay among the lipidome, anthracyclin uptake, ER stress, and sensitivity to anthracyclins

The metabolome, the lipidome, the proteome, and the transcriptome affect anthracyclin uptake and ER stress^58,59^. Anthracyclins themselves affect the ‘omes, which further modifies the uptake of the drug and ER stress. The outcome of this crossed interplay is the different mechanisms of cell death and hence the sensitivity of a cell/tumor type to anthracyclins^13^. Of special interest in this interplay is the lipidome, as it conditions: i) drug uptake (drug-membrane interaction and membrane fluidity), ii) ER stress (ratio PC/PE, balance of PUFAs versus cholesterol and saturated lipids), and iii) the type of cell death (lipids with PUFAs in ferroptosis) (Scheme 3).

Regarding anthracyclin uptake, lipid membrane composition determines its fluidity and drug membrane permeation, as well as activation of drug carrier-mediated efflux mechanisms related to multidrug resistance^9,60^. We have found that the intracellular uptake ratio of DOX was one order of magnitude higher in HepG2 or Huh7 than in SNU449 (HepG2 > Huh7 >> SNU449, Kullenberg, et al., submitted^20^). Consequently, we speculate that the different response of the lipidome of the three cell lines (Figure 3) may play a role in the higher uptake of DOX by HepG2 and Huh7. Interestingly, PGs presented a remarkable increase in HepG2 and Huh7 with both anthracyclins (Figure 3). PGs are negatively charged, and interact in the membrane with positively charged DOX and IDA. Other lipids also affect the properties of the membranes regarding drug uptake: i) the levels of other negatively charged lipids (PIs, PSs); ii) zwitterionic lipids (PCs, PEs)^10^; and fluidity (cholesterol and saturation of fatty acids in the glycerolipids). Our study was limited to three cell lines, which prevents the discussion of these multiple changes. Nevertheless, in the light of previous research, we speculate that the strong enrichment of PGs in the membranes of HepG2 and Huh7 (Figure 3) may play a role in favoring anthracyclin uptake.

ER stress plays an essential role in apoptosis and ferroptosis^61^, as well as in promoting tumorigenesis in HCC^62,63^. In addition, ER stress mediates drug resistance to chemotherapeutic agents^64^. There is a crosstalk between ER stress pathways and the lipidome^65^. In this crosstalk, more than the composition in a specific type of lipid, it is the balance among different types of lipids what is associated with ER stress. For example, the imbalance between lipids with PUFAs versus saturated and cholesterol provokes ER stress^58,59^. In this context, we observed that: i) membrane lipids with PUFAs increased in general in each of the three cell lines (Section 4.3), ii) membrane lipids with saturated and mono-unsaturated fatty acids did not present any clear trend (Section 4.2), and ii) cholesterol lipids increased in Huh7 and SNU449, but not in HepG2 (Figure 3). This ensemble of observations suggests a higher imbalance in HepG2 cells than in SNU449 (Scheme 3), which in the light of previous research partially explains the stronger increase of ER stress in HepG2 cell than in SNU449 (Figure 8). Many other factors beyond the scope of this study affect ER stress ^65^, which may explain why Huh7 did not present a strong imbalance between lipids with PUFAs and sterols but presented a strong increase of ER stress (Figure 8B).

**Scheme 3.**
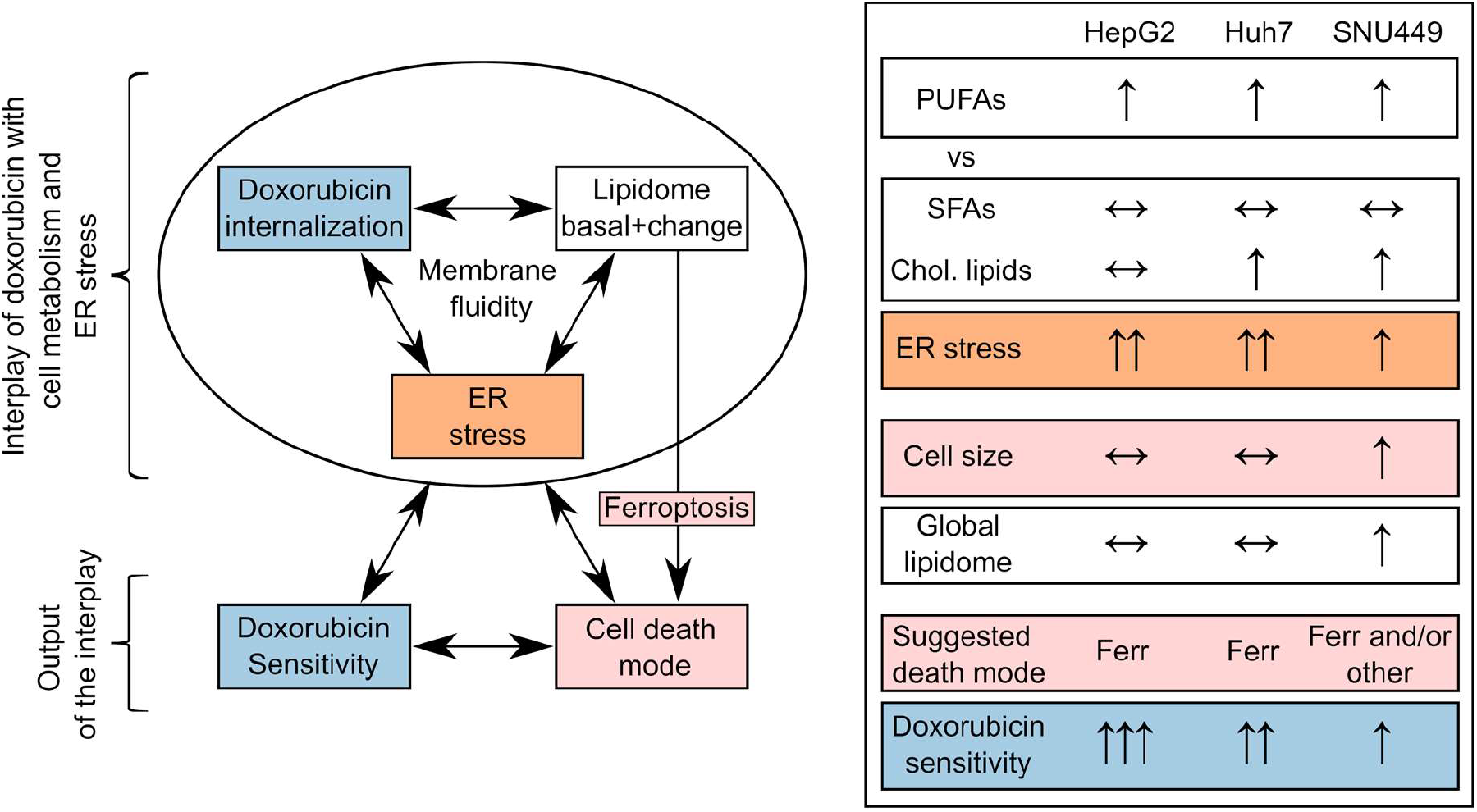
On the left, schematic interplay between DOX, the lipidome, and ER stress for a cell type, with the output of sensitivity to the drug and cell death mode. On the right, ensemble of qualitative trends of DOX treatment for the three cell lines: i) the balance between PUFAs lipids versus saturated fatty acids (SFAs) and cholesterol lipids (Chol. Lipids), ii) ER stress, iii) cell size and the global lipidome, iv) suggested death mode (Ferr for ferroptosis) and DOX sensitivity.

Considering the ensemble data about the lipidome, drug uptake, and ER stress (Scheme 3), the observed changes may contribute to explain why SNU449 is one of the most resistant cell lines to anthracyclins^66^. In comparison with HepG2 and Huh7, SNU449 presented: i) an increase in size (Figure 2), ii) a general increase in the lipidome (Figure 3), iii) a weaker increase in ER stress makers (Figure 8). Despite other factors (metabolome, proteome, transcriptome) also take part in the cell death, this differential behavior in the context of previous research suggests a key role of lipids in cell death. In this role, more than the change in one lipid or lipid family, it is the ensemble of lipids which seems involved in the resistance to anthracyclins.

In conclusion, as discussed in previous paragraphs, the sensitivity to anthracyclins can be related to a complex interplay among the lipidome, cellular uptake and intracellular disposition of drugs, ER stress, and the mechanism(s) of cell death. These factors depend on one another and on other factors beyond the scope of this study (transcriptome, proteome, metabolome). This interplay prevents the discussion of the effect of a lipid or a family of lipids in isolation. Despite these limitations, previous research give sense to the observed changes in the lipidome in relation to drug uptake, ER stress, and the sensitivity of the cells to anthracyclins for HepG2, Huh7, and SNU449.

### 4.6. Perspectives in metabochemotherapy and anthracyclins

It is known that PUFAs potentiate the effect of DOX in vitro and in animal models^30,67^. This effect might be general, as it would potentiate the lipid peroxidation in ferroptosis induced by anthracyclins. However, potentiating ferroptosis may increase the cardiotoxicity of anthracyclins^68^. Paradoxically, it has been reported that PUFAs may have a cardioprotective effect in anthracyclin chemotherapy^69,70^. This suggests that PUFAs may present a therapeutic window. In this window, PUFAs may potentiate anthracyclin-induced ferroptosis in tumor cells, and in parallel minimizing cardiotoxicity. Similarly, the administration of anthracyclins in bilayers with etherPEs with PUFAs may also present a therapeutic window. Our results suggests that this would be the case for tumors with cells with lipid metabolism similar to HepG2 and Huh7. As discussed before, ferroptosis might not be the only suggested mechanism of death for SNU449. Nevertheless, we speculate that the treatment with PUFAs and/or etherPEs with PUFAs might tumors with cells like SNU449 into the path of lipid peroxidation and ferroptosis. In this perspective, PUFAs and etherPEs with PUFAs could be considered general adjuvants of anthracyclins.

Another suggested adjuvant of anthracyclins is the inhibition of cholesterol formation by statins^71^. Our results suggest that the effect of anthracyclins on cholesterol lipids was specific for each of the the three cell lines of the same type of tumor. The fact of having a different behavior for three cell lines of the same type of tumor indicates that the modulation of the cholesterol pathway would be of interest depending on both the cancer type and the patient. This agrees with previous literature, as statins present synergizing and protective effects in different models^29,72–74^. Furthermore, the adjuvant effect of statins is not clear in patients^71^. An explanation would be that the adjuvant effect of statins might depend on the patient pharmacokinetics, pharmacodynamics, and cholesterol metabolism. Consequently, while statins do not seem to be a general adjuvant of anthracyclins, it seems plausible that they could be used for specific tumors and in personalized metabochemotherapy.

## 5. Conclusions

To the best of our knowledge, here we present for the first time that the upregulation of free PUFAs and etherPEs with PUFAs is a general trait of anthracyclins. According to literature, this increase is pro-ferroptotic by promoting lipid peroxidation but other metabolic changes might counteract this type of cell death. This is the case of cholesterol lipids and saturated and monounsaturated glycerolipids, which are markers of the de novo lipogenesis in the LXR/SREBP pathways. PGs, which are negatively charged at physiological and pathophysiological pH, presented a strong increase in HepG2 and Huh7. This increase may play a key role in the higher uptake ratio of DOX of this cell lines. We found that these lipids showed disparate behavior depending on the cell type after the treatment with anthracyclins. These specificities partially explain the different sensitivity to anthracyclins by the effect of lipids on passive diffusion of the drugs, their metabolites, membrane fluidity, ER stress, and mechanism(s) of cell death.

In conclusion, the behavior of the lipidome in the three cell lines suggests that: i) PUFAs or PUFA-containing lipids may potentiate the effect of anthracyclins in a general way, and ii) statins may potentiate or attenuate the effect of anthracyclins in a cancer/patient specific way. Further research in animal models and patients is warranted to establish the possibility of transferring these observations from bench to bedside.

## Acknowledgments

We are grateful to Dilruba Ahmed (Karolinska Institute, Sweden) for supplying Huh7 cell line.

